# Episodic Positive Selection Signatures in *Arabidopsis* CONSTANS-like Genes COL3 and COL5 Indicating Adaptive Evolution in Red-Light Signaling Pathways

**DOI:** 10.1101/2025.04.03.646976

**Authors:** Krishnendu Sinha

## Abstract

This study investigates the adaptive evolution of the CONSTANS-like (COL) gene family in *Arabidopsis thaliana* (L) Heynh., focusing on the molecular and structural consequences of episodic positive selection. Using CODEML-based site and branch-site models, I identified statistically significant adaptive substitutions in COL3 and COL5. Ancestral sequence reconstruction revealed that COL3 evolved from serine to alanine at residue 212, while COL5 transitioned from glycine to serine at residue 343. To assess structural impacts, I retrieved modern COL protein structures from the AlphaFold Protein Structure Database, introduced reverse mutations, and generated ancestral models using AlphaFold3. Stability analyses with DynaMut indicated that the G343S mutation in COL5, located within a unique proteomic domain, resulted in slight destabilization and reduced flexibility, with ChimeraX revealing the loss of hydrogen bonding. In contrast, the S212A mutation in COL3, positioned upstream of its CCT domain, showed a stabilizing effect despite reduced flexibility. GO enrichment analysis revealed that positively selected COL genes are significantly associated with red and far-red light signaling and developmental regulation. These findings suggest that episodic positive selection fine-tunes COL protein function, warranting further studies using molecular dynamics simulations and in vivo functional assays.

## 1. Introduction

Plants adjust a wide range of physiological and developmental processes in response to the natural cycle of light and darkness. Photoperiodism, defined as the ability to sense and react to changes in daylength, plays a critical role in timing key events such as flowering, bud dormancy, and tuber formation across different species^1^. Since the duration of daylight varies with season and latitude, plants rely on internal timing mechanisms, including a circadian clock, to accurately measure these changes despite environmental fluctuations^1^. This precise temporal regulation allows plants to coordinate growth and reproduction with favorable seasonal conditions and to optimize responses to abiotic stresses such as cold, drought, and salinity^2^. Additionally, photoperiodic cues can modulate defense responses, influencing interactions with pathogens and other environmental challenges^2^. Overall, the integration of external light signals with intrinsic developmental programs via photoperiodism is essential for maximizing resource use, ensuring reproductive success, and enhancing stress tolerance in a wide variety of plant species.

The CONSTANS-like (CO-like; COL) genes regulate the precise timing of flowering by integrating internal circadian rhythms with external light cues, a process that is critical for plant reproductive success under varying environmental conditions^3^. Flowering is a critical trait that determines reproductive success, and its precise timing is essential for successful pollination, seed development, and environmental adaptation^3^. In *Arabidopsis thaliana* (L) Heynh. (hereafter *Arabidopsis*), the COL gene serves as the central integrator of photoperiodic flowering pathways, acting as a nuclear zinc finger transcription factor characterized by two N-terminal B-box domains and a C-terminal CCT domain^4^. *Arabidopsis* possesses 17 COL genes that are phylogenetically subdivided into three groups, with Group I (including CO and COL1–COL5) containing an additional conserved six–amino acid motif at the C-terminus^4,5^. CO expression, regulated by the circadian clock component GIGANTEA (GI), is confined to the vascular tissue of leaves, where it plays a pivotal role in mediating xylem expansion and stomatal opening while integrating internal rhythms with external day-night cycles^6^. Under long-day conditions, CO promotes the transcription of key flowering regulators such as FLOWERING LOCUS T (FT), SOC1, and TSF, thereby initiating the flowering process^4^. This CO/FT module, though highly conserved among photoperiod-sensitive plants, exhibits functional variation across species, as seen with rice Hd1, suggesting both conserved and divergent evolutionary adaptations^4,5^. Moreover, beyond flowering time regulation, COL genes in *Arabidopsis* display diverse roles, including the regulation of photomorphogenesis, root development, shoot branching, and responses to abiotic stresses^4,5^. These overlapping functions underscore the complexity and evolutionary significance of the COL gene family in *Arabidopsis*, providing a robust model for understanding the molecular mechanisms underlying plant development and adaptation. In *Arabidopsis*, despite their established roles in modulating several pathways, the evolutionary forces shaping the COL gene family have not been comprehensively characterized.

The diversification of gene families is fundamental to plant evolution, enabling species to develop new functions and adapt to varying environments. In this context, the COL gene family, which is central to photoperiodic flowering and light signal transduction, serves as an excellent model for investigating molecular evolution and the evolution of photoperiodism^3^. By employing rigorous sequence quality control, detailed phylogenetic reconstruction, codon-based selection analyses, and comprehensive structural modeling, this study uncovers adaptive modifications that have fine-tuned COL proteins over evolutionary time. This multifaceted approach enables the detection of specific amino acid substitutions favored by natural selection and permits inference of their structural and functional consequences—such as alterations in protein–protein interactions and stability that may ultimately affect photoperiodic responses. These findings shed light on how selection pressures shape regulatory networks at the molecular level, providing a robust framework for understanding the evolution of photoperiodism as a key adaptive trait in plants.

## 2. Materials and Methods

### 2.1. Gene Identification and Sequence Retrieval

COL gene identifiers were initially obtained from the Plant Transcription Factor Database (PlantTFDB, v5.0) and further cross-validated using The Arabidopsis Information Resource (TAIR) to ensure comprehensive gene annotation. Coding sequences (CDS) corresponding to the identified COL genes were then downloaded from the National Center for Biotechnology Information (NCBI). When multiple transcript variants were available for a given gene, the longest transcript was selected to represent the gene^7^. Total 17 CDSs were retrieved in this process.

### 2.2. Sequence Quality Control

The retrieved CDS were subjected to a rigorous quality control protocol implemented via a custom Python script^7^. Each sequence was checked for the presence of a canonical start codon (ATG) and an in-frame stop codon at the 3′ end, and any sequences containing premature or internal stop codons were discarded. In addition, sequences were filtered to retain only those with lengths divisible by three and with a minimum length threshold of 300 nucleotides. Outliers with unusually long or short CDS, which could negatively impact downstream alignment, were excluded based on interquartile range criteria. The script used for quality control All filtering parameters and thresholds are documented in Supplementary File 1. Finally, all 17 sequenced passed the quality check and selected for further study (Table 1).

**Table 1.**
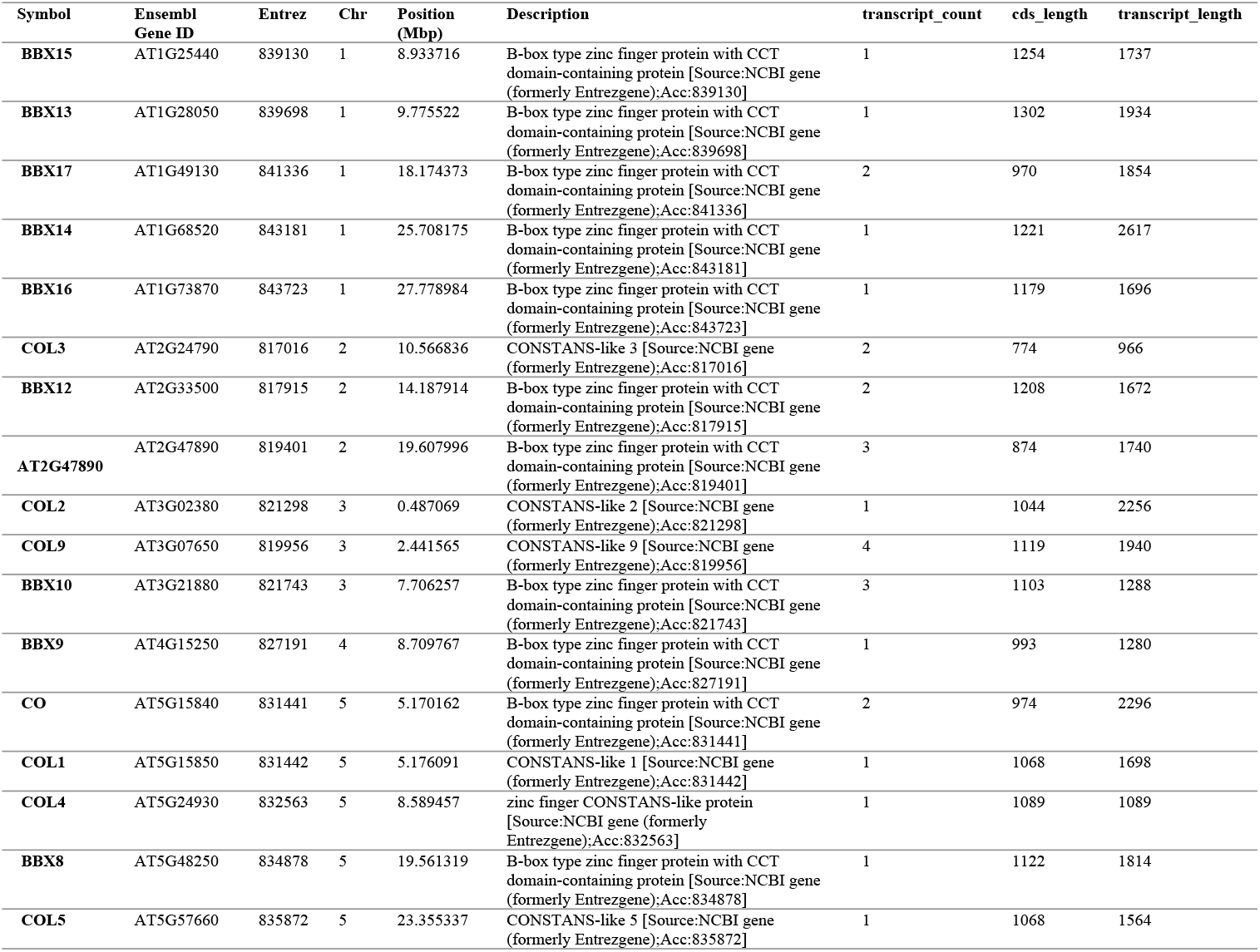
Detailed description of 17 *Arabidopsis* COL CDSs selected for the study.

### 2.3. Multiple Sequence Alignment and Phylogenetic Reconstruction

High-quality COL CDS were aligned using the PRANK algorithm (version 170427) with default parameters optimized for codon alignments^7,8^. The resulting multiple sequence alignment (MSA) was manually inspected for consistency using Aliview (v1.30)^9^. A maximum likelihood (ML) phylogenetic tree was subsequently inferred with IQ-TREE (multicore version 2.0.7) using the best-fit codon substitution model SCHN05+F+R3 chosen according to BIC^10,11^. The tree was constructed with 10,000 ultrafast bootstrap replications to assess node support. The complete IQ-TREE command line was iqtree2 -s msa.fas -alrt 10000 -B 10000 -st CODON -T AUTO -bnni. MSA file and IQ-TREE2 log file are provided as Supplementary File 2 and 3, respectively.

### 2.4. Codon-Based Selection Analysis and Ancestral Sequence Reconstruction

To detect signatures of positive selection, codon-based analyses were performed using CODEML from the PAML package (v4.9j)^12^. Multiple site models (M0, M1a, M2a, M7, and M8) were tested using the MSA and phylogenetic tree generated earlier^13^. Likelihood ratio tests (LRTs) were used to compare nested models (e.g., M1a vs. M2a and M7 vs. M8)^13^. For the branch and branch-site models, a custom Python script was used to sequentially assign each branch, including ancestral nodes, as the foreground, and these assignments were analyzed concurrently^7^. To adjust for multiple testing across branches, the Benjamini–Hochberg procedure was applied to the resulting p-values, with adjusted p-values below 0.05 considered statistically significant^14^. The ω (dN/dS) values for the modern and ancestral genes were derived from the branch model output and then compared with those of the modern genes using a Mann-Whitney U test, with significance set at p < 0.05^15,16^. Ancestral sequence reconstruction was performed using the free-ratio model in CODEML with the RateAncestor option set to 1^17^. Codeml control file, ASR control file and ASR output logs, are provided in Supplementary Files 4, 5 and 6, respectively.

### 2.5. Gene Ontology Enrichment Analysis

To assess the functional implications of adaptive changes in COL genes, I performed Gene Ontology (GO) enrichment analysis using the ShinyGO web server (v0.82) ^18^. The analysis was conducted on two datasets: one comprising genes exhibiting episodic positive selection and the other containing the remaining COL genes. Focusing on the Biological Process category, enrichment significance was determined using a hypergeometric test ^18^. P-values were then adjusted for multiple comparisons using the default false discovery rate (FDR) settings, and GO terms with an FDR-adjusted p-value below 0.05 were considered significantly enriched.

### 2.6. Structural Modeling and Protein Stability Predictions

Tertiary protein structures for COL3 and COL5 were retrieved from the AlphaFold Protein Structure Database^19^. For structural evaluation of adaptive mutations at the selective sites, I obtained the corresponding modern COL protein sequences from NCBI and introduced specific mutations at the positively selected sites, as identified by CODEML-based ancestral sequence reconstruction. The analysis indicated that in COL3 the ancestral state at residue 212 was serine while the modern protein has alanine, and in COL5 the ancestral state at residue 343 was glycine compared to a serine in the modern protein. To assess the impact of these adaptive changes, I introduced the mutations A212S for COL3 and S343G for COL5, thereby generating mutation-derived protein sequences. These modified sequences were modeled using the AlphaFold3 web server, and the resulting structures were exported in PDB format^20^. All the tertiary structures were further analyzed using ChimeraX (v1.9) to visualize differences in domain organization and hydrogen-bond networks^21^. To quantify the impact of the substitutions on protein conformation, stability, and flexibility, I used DynaMut^22^, which computes changes in free energy (ΔΔG) and vibrational entropy (ΔΔSVib) by specifying the mutations S212A and G343S. This approach provides a reproducible framework for understanding the structural consequences of adaptive evolution in the COL gene family.

### 2.7. Statistical Analyses

All statistical tests were performed using R (v4.4.3) in the RStudio environment and Python. For selection analyses, LRT statistics obtained from CODEML using Supplementary File 7, were compared against the chi-square distribution with the appropriate degrees of freedom, and p-values were adjusted using the Benjamini–Hochberg method to control the false discovery rate when analyzing multiple foreground branches in branch and branch-site models using custom Python scripts (Supplementary File 8). The Mann-Whitney U test in R (Supplementary File 9) was used to compare ω values between ancestral and modern sequences. In all analyses, statistical significance was defined as p < 0.05 unless stated otherwise.

## 3. Results

### 3.1. Seventeen Sequences Passed Quality Checks and the ML Tree Shows Three Distinct Clades

All 17 COL CDSs were retained after rigorous quality control resulting in a refined dataset that provided a reliable basis for downstream analyses. The COL sequences were aligned using the PRANK algorithm, and the resulting multiple sequence alignment was of high quality checked through AliView. IQTREE2 produced a robust ML tree supported by 10,000 bootstrap replicates, revealed 3 distinct clades within the COL family, indicating that the gene family has undergone diversification (Figure 1). The phylogenetic clusters corresponded well with previously reported subgroups^4^, supporting the notion that different COL paralogs may have evolved specialized roles in photoperiodic regulation.

**Figure 1.**
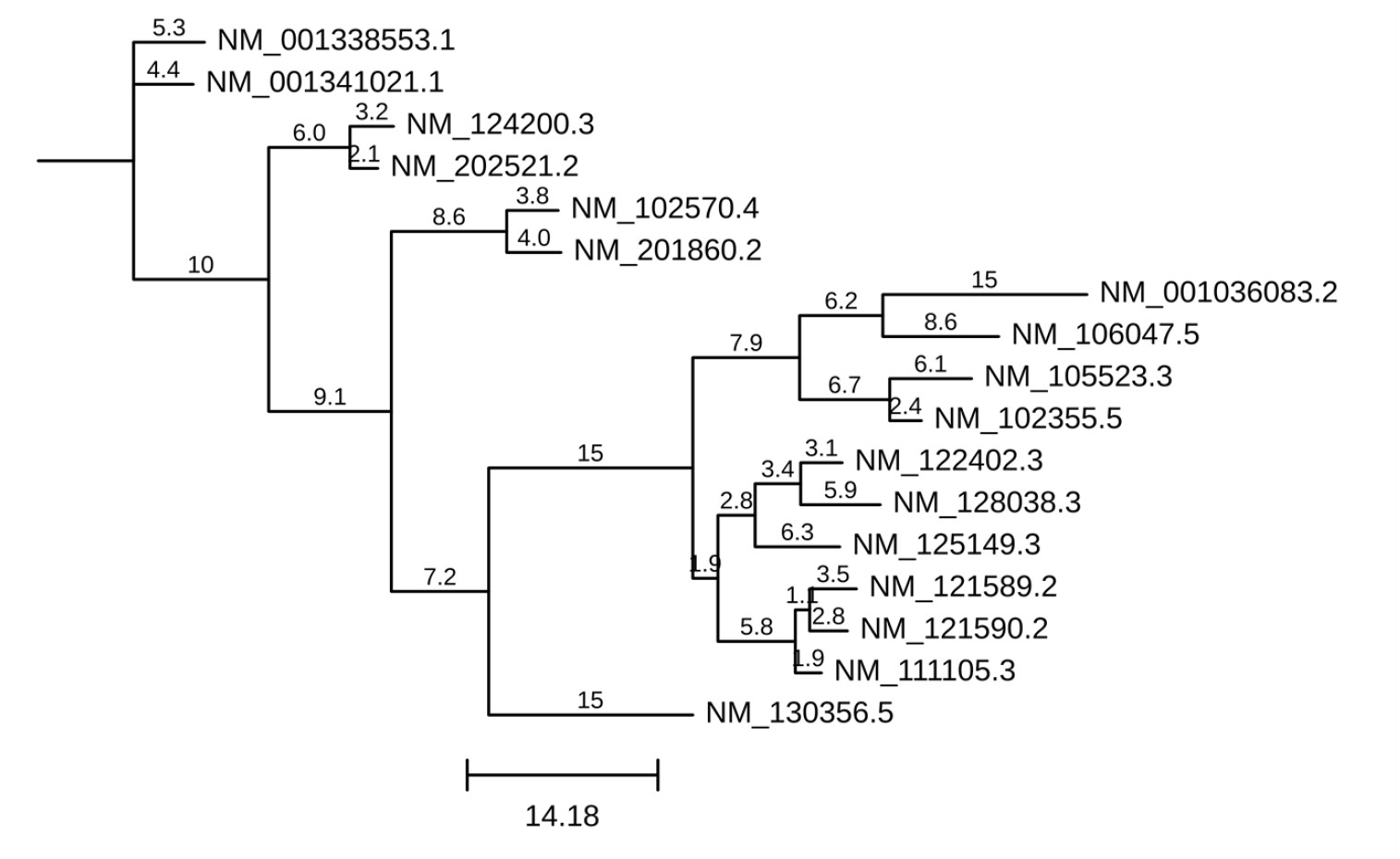
Phylogenetic tree for *Arabidopsis* COL gene family reconstructed by maximum-likelihood (ML) under the SCHN05+F+R3 model with IQ-TREE multicore v2.0.7 using the CDS. The Tree was visualized using TreeViewer v2.2.0^24^. The tree is arbitrarily rooted for better visualization and the numbers showing branch lengths.

### 3.2. CODEML Analysis Spots Episodic Selection Signature in Three Genes

To assess selective forces acting on the *Arabidopsis* COL gene family, I first applied CODEML site models to evaluate variability in ω (dN/dS) ratios across codon sites. Comparisons between the M0 and M1a models revealed significant variation in selective pressures (LRT: 2Δℓ = 264, df = 1, p < 1e−4), indicating that while some codon sites are under purifying selection (ω < 1) and others evolve neutrally (ω ≈ 1). No evidence of positive selection was detected with these models. Subsequent comparisons using the M1a versus M2a and M7 versus M8 models also failed to reveal statistically significant signals of positive selection (LRT: 2Δℓ = 0 and 2Δℓ = 0.1, df = 2; p > 0.05, respectively). These findings suggest a predominance of purifying selection with certain sites evolving neutrally (Supplementary File 10).

Then statistically more prominent free-ratio branch model was employed to investigate lineage-specific selection by estimating ω values for individual branches. This analysis identified two branches (NM_121589.2 and NM_121590.2) with ω ratios of 0.92 and 0.44, respectively, which were statistically higher than the background ω of 0.22; however, these values remained below the threshold of 1, indicating that the signal of positive selection was not robust (Supplementary File 10).

To further refine the analysis and found episodic selection signature, the statistically most robust model, branch-site models (A and A1)^7,12^ was applied, designating each branch as the foreground in turn. This stringent approach detected significant episodic positive selection on two foreground branches, NM_125149.3 (COL5), and NM_128038.3 (COL3), with LRTs of 2Δℓ = 8.06 and 11.58 (df = 1, p < 0.05), respectively (Table 2, 3 and 4). Branch-site models further supported episodic positive selection for NM_125149.3 and NM_128038.3, with significantly elevated ω values (p < 0.01) and Bayes Empirical Bayes (BEB) analysis identifying positively selected sites corresponding to residues 1764 (COL5) and 1505 (COL3) in the multiple sequence alignment, which align to positions 343 and 212 in the ungapped sequences. This analysis showed, significant heterogeneity in selective pressures was observed across the *Arabidopsis* COL gene family, with robust evidence of episodic positive selection detected in COL3 and COL5. These findings imply that episodic positive selection may have contributed to the functional diversification and alterations in structural stability and flexibility within the COL gene family.

**Table 2.** Result of BH correction on LRT test result for Branch-Site (BS) model. Note: - BS_NULL stands for null model.

| Gene | lnL_BS | lnL_BS_NULL | LRT | p_value | FDR_Corrected_P | Significant (FDR < 0.05) |
| --- | --- | --- | --- | --- | --- | --- |
| **NM_001036083.2** | -18262.79417 | -18263.60718 | 1.626022 | 0.202254146 | 0.42979006 | No |
| NM_001338553.1 | -18260.81343 | -18263.01051 | 4.394158 | 0.036062263 | 0.102176411 | No |
| NM_001341021.1 | -18263.64263 | -18263.64263 | 0 | 1 | 1 | No |
| NM_102355.5 | -18265.38787 | -18265.38788 | 2E-06 | 0.998871621 | 1 | No |
| NM_102570.4 | -18263.39947 | -18263.87034 | 0.941748 | 0.331828706 | 0.5641088 | No |
| NM_105523.3 | -18264.82763 | -18265.32542 | 0.995586 | 0.318380929 | 0.5641088 | No |
| NM_106047.5 | -18261.82293 | -18264.44239 | 5.238918 | 0.022087007 | 0.102176411 | No |
| NM_111105.3 | -18265.38787 | -18265.38787 | 0 | 1 | 1 | No |
| NM_121589.2 | -18262.66066 | -18262.66066 | 0 | 1 | 1 | No |
| NM_121590.2 | -18265.12387 | -18265.12387 | 0 | 1 | 1 | No |
| NM_122402.3 | -18262.54828 | -18264.9048 | 4.71305 | 0.029934498 | 0.102176411 | No |
| NM_124200.3 | -18265.38788 | -18265.38787 | -2E-06 | 1 | 1 | No |
| NM_125149.3 | -18257.20117 | -18261.23555 | 8.068754 | 0.004503507 | 0.038279813 | Yes |
| NM_128038.3 | -18258.53121 | -18264.32234 | 11.58227<br>8 | 0.000665833 | 0.011319165 | Yes |
| NM_130356.5 | -18261.44656 | -18262.30741 | 1.721704 | 0.18947385 | 0.42979006 | No |
| NM_201860.2 | -18260.61796 | -18262.94535 | 4.65479 | 0.03096702 | 0.102176411 | No |
| NM_202521.2 | -18265.38787 | -18265.38787 | 0 | 1 | 1 | No |

**Table 3.** Maximum-likelihood Estimates of Parameters in the ω Distribution Under the Branch-site Model A. Note.-The branch-site model assumes four site classes (0, 1, 2a, 2b), with different ω ratios for the foreground and background lineages. Sites from site class 0 are under purifying selection along all branches with 0 < ω0 < 1, while all branches in site class 1 are undergoing neutral evolution with ω1 = 1. In site classes 2a and b, there is positive selection along foreground branches with ω2 > 1, while the background branches are under purifying selection with 0 < ω0 < 1 or undergoing neutral evolution with ω1 = 1.

| Foreground branch | Site class | Proportion | Background $\omega$ | Foreground $\omega$ |
| --- | --- | --- | --- | --- |
| NM_106047.5 | 0 | p0 = 0.56778 | $\omega_0 = 0.14180$ | $\omega_0 = 0.14180$ |
| | 1 | p1 = 0.35242 | $\omega_1 = 1.00000$ | $\omega_1 = 1.00000$ |
| | 2a | p2a = 0.04923 | $\omega_0 = 0.14180$ | $\omega_2 = 393.04443$ |
| | 2b | p2b = 0.03056 | $\omega_1 = 1.00000$ | $\omega_2 = 393.04443$ |
| NM_125149.3 | 0 | p0 = 0.56728 | $\omega_0 = 0.14082$ | $\omega_0 = 0.14082$ |
| | 1 | p1 = 0.34500 | $\omega_1 = 1.00000$ | $\omega_1 = 1.00000$ |
| | 2a | p2a = 0.05455 | $\omega_0 = 0.14082$ | $\omega_2 = 7.90150$ |
| | 2b | p2b = 0.03317 | $\omega_1 = 1.00000$ | $\omega_2 = 7.90150$ |
| NM_128038.3 | 0 | p0 = 0.58403 | $\omega_0 = 0.14313$ | $\omega_0 = 0.14313$ |
| | 1 | p1 = 0.35913 | $\omega_1 = 1.00000$ | $\omega_1 = 1.00000$ |
| | 2a | p2a = 0.03520 | $\omega_0 = 0.14313$ | $\omega_2 = 132.40748$ |
| | 2b | p2b = 0.02165 | $\omega_1 = 1.00000$ | $\omega_2 = 132.40748$ |

**Table 4.** Branch-site Test of Positive Selection Along Three Foreground Branches. Note.—The critical values are χ21,5%=3.84 (i.e., 1 degree of freedom, df, at 5% significance) and, χ21,1%=6.63 (i.e., 1 degree of freedom, df, at 1% significance). The LRT statistic, 2Δℓ, is reported for all model comparisons. Sites that Have a Probability Higher Than 95% to be Under Positive Selection According to the BEB Analysis Under the Branch-site Model A.

| Model | $\ell$ | Free parameters<sup>a</sup> | df | $2\Delta\ell$ | $\text{Pr}(\omega > 1) > 95\%$ |
| --- | --- | --- | --- | --- | --- |
| NM_106047.5 as the foreground (unrooted) |  |  |  |  |  |
| Null model (model A with $\omega_2 = 1$) | $\ell_0 = -$<br>18264.442387 | 38 | 1 | 5.238918 | |
| Alternative model (model A) | $\ell_1 = -$<br>18261.822928 | 39 | | | |
| NM_125149.3 as the foreground (unrooted) |  |  |  |  |  |
| Null model (model A with $\omega_2 = 1$) | $\ell_0 = -$<br>18261.235545 | 38 | 1 | 8.068754 | 1764 |
| Alternative model (model A) | $\ell_1 = -$<br>18257.201168 | 39 | | | |
| NM_128038.3 as the foreground (unrooted) |  |  |  |  |  |
| Null model (model A with $\omega_2 = 1$) | $\ell_0 = -$<br>18264.322344 | 38 | 1 | 11.58228 | 1505 |
| Alternative model (model A) | $\ell_1 = -$<br>18258.531205 | 39 | | | |

### 3.3. Ancestral Sequence Reconstruction Confirms Episodic Selection and Dilution of Selection Pressure in Modern Genes

Ancestral sequence reconstruction using CODEML’s site model enabled the inference of the most probable ancestral amino acid states at key positions. For both COL3 and COL5, the ancestral sequences differed from the modern proteins at the positively selected sites, providing further evidence that episodic positive selection has shaped these proteins over evolutionary time. Additionally, a comparison of the ω values between the reconstructed ancestral sequences and modern sequences, as determined by the Mann-Whitney U test, revealed statistically significant differences (p < 0.05) (Figure 2). This finding implies that the overall intensity of purifying selection has diluted in modern genes relative to their ancestral counterparts, consistent with observations reported for GmWRKY genes^7^.

**Figure 2.**
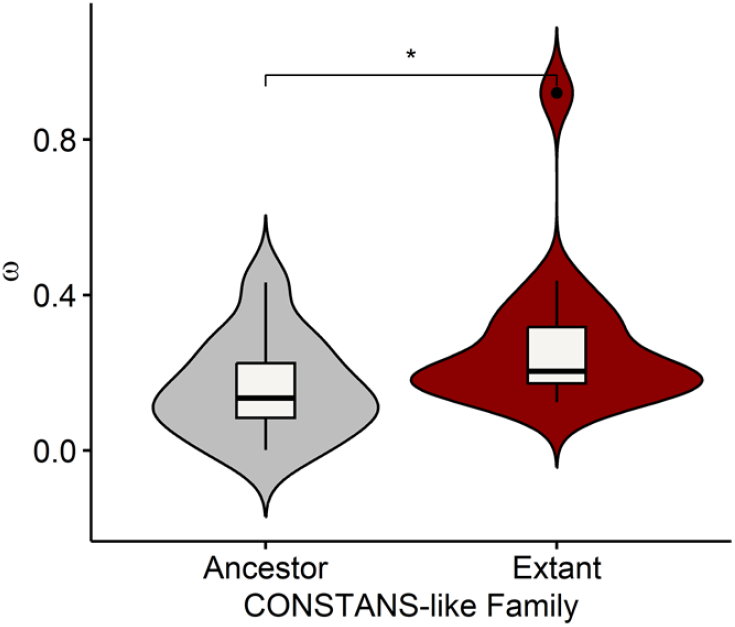
Violin plots comparing ancestral and modern *Arabidopsis* COL genes based on Mann–Whitney U tests for omega (ω) ratio (dN/dS). Significant differences (*p < 0.05) between ancestral and extant sequences were observed. The width of each violin represents the distribution of values and horizontal lines indicate the median.

### 3.4. Gene Ontology Enrichment Analysis

Functional categorization of the positively selected COL genes was performed using GO enrichment analysis with ShinyGO. The analysis was conducted separately on two groups: one group comprised two COL genes that exhibited clear episodic positive selection signatures, while the other contained the remaining 15 COL genes. The results revealed that the two genes under episodic positive selection were significantly enriched (BH-adjusted p < 0.05) for biological processes associated with red and far-red light signaling (Figure 3a). In contrast, the other group did not display significant enrichment for these pathways (Figure 3b). These findings suggest that positive selection favors COL genes involved in red/far-red light perception and key developmental processes, highlighting their potential adaptive role in modulating plant growth and reproductive timing.

**Figure 3.**
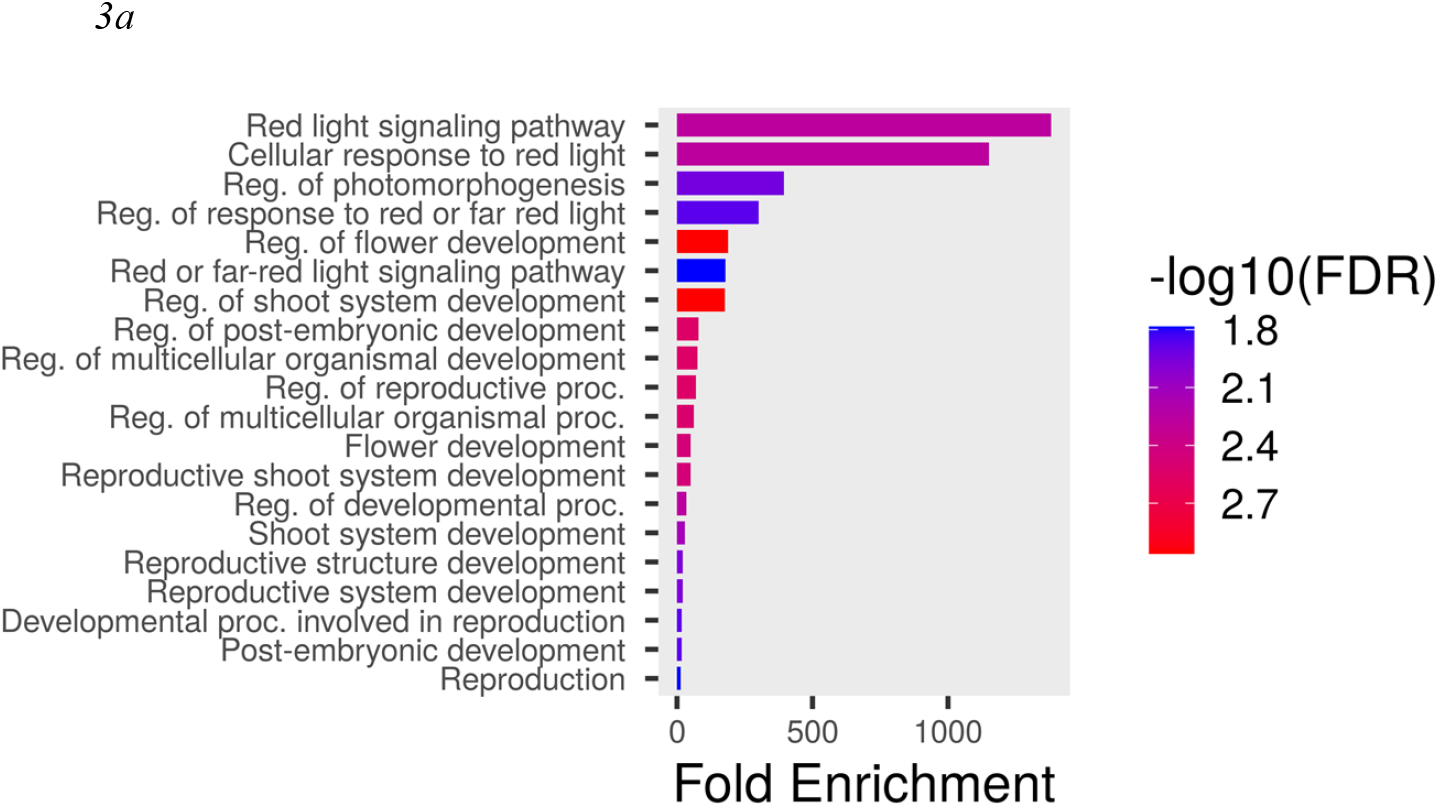
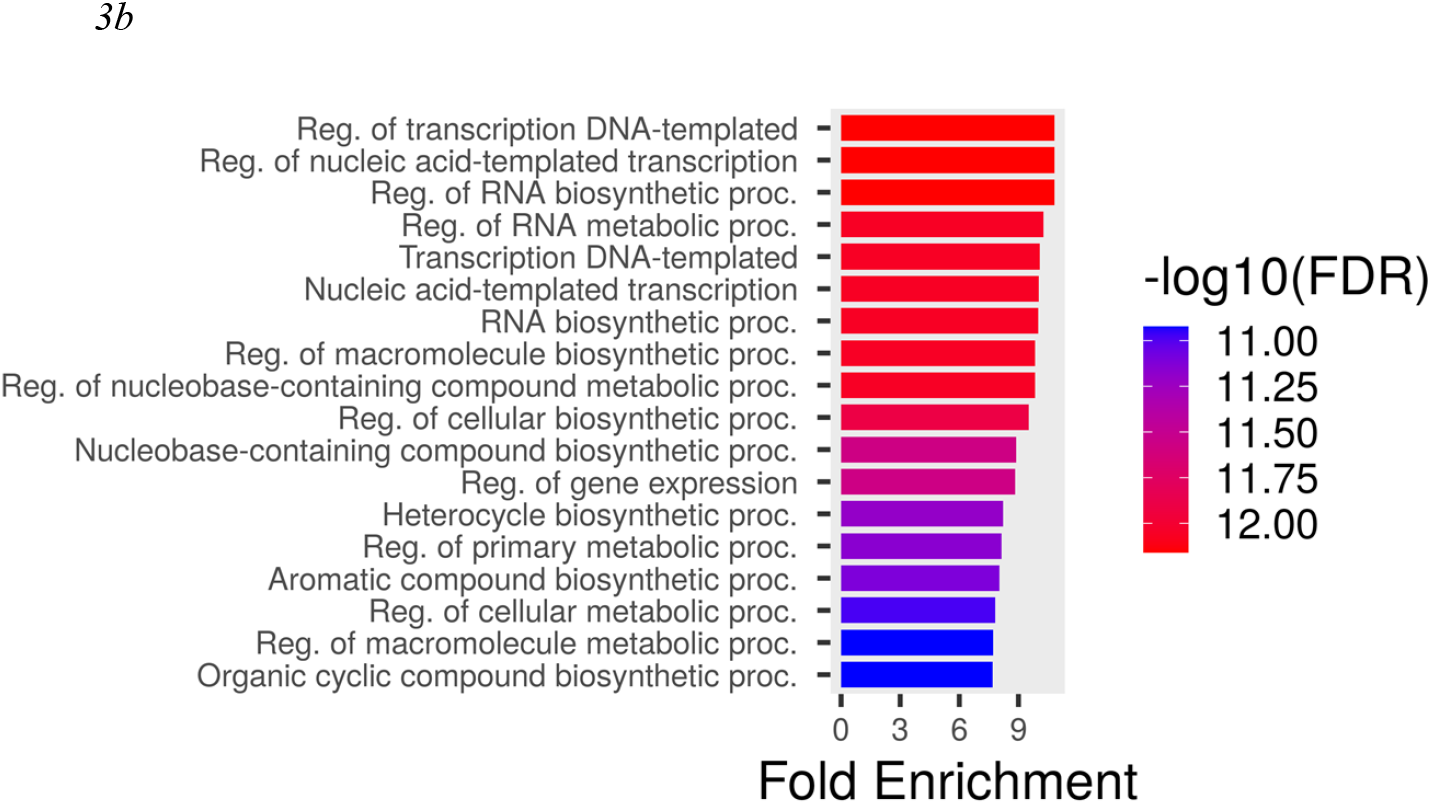
GO enrichment analysis plot. (a) Enrichment plot using group of episodic positively selected genes (Supplementary File 11). (b) Enrichment plot using group of rest of the genes (Supplementary File 12).

### 3.5. Structural Modeling Shows Enhanced Stability with Adaptive Mutations

To assess the impact of adaptive mutations on protein structure, modern COL protein structures retrieved from the AlphaFold Protein Structure Database were visualized using ChimeraX to detect any conformational changes. In addition, DynaMut was used to predict changes in protein stability and flexibility. For COL5, the mutation at residue 343 (G343S) is particularly notable because this residue lies within a unique proteomic domain outside the canonical CCT domain (which spans residues 285–327)^23^. DynaMut predicted a ΔΔG of 0.195 kcal/mol, indicating a slight stabilizing effect^22^. Additionally, the ΔΔSVib value from ENCoM (–0.047 kcal·mol^−1^·K^−1^) indicates a reduction in protein flexibility^22^. Structural inspection using ChimeraX revealed that the modern COL5 protein exhibits hydrogen bonds involving the serine at position 343, bonds that are absent when S is replaced by G (Figure 4a, 4b). In contrast, for COL3, the adaptive mutation occurs at residue 212 (S212A), which is located just upstream of the CCT domain (spanning residues 229–271)^23^. For COL3, DynaMut predicted a ΔΔG of 0.735 kcal/mol, suggesting an overall stabilizing effect where as a larger reduction in flexibility was observed (ΔΔSVib ENCoM: –0.162 kcal·mol^−1^·K^−1^)^22^. Notably, no significant differences in hydrogen bonding were observed in COL3 between the wild-type and mutant forms (Figure 4d, 4e). But in for both the protein structural modelling shows significant conformational changes due to adaptive mutations (Figure 4c, 4f). Together, these findings suggest that the G343S mutation in COL5 likely increase key stabilizing interactions and reduces structural flexibility, whereas the S212A mutation in COL3 may enhance stability while altering flexibility. This differential impact of adaptive mutations highlights the nuanced role of episodic positive selection in fine-tuning the functional dynamics of COL proteins, conformation of which warrant more in-depth studies using protein-protein and protein-DNA interactions.

**Figure 4.**
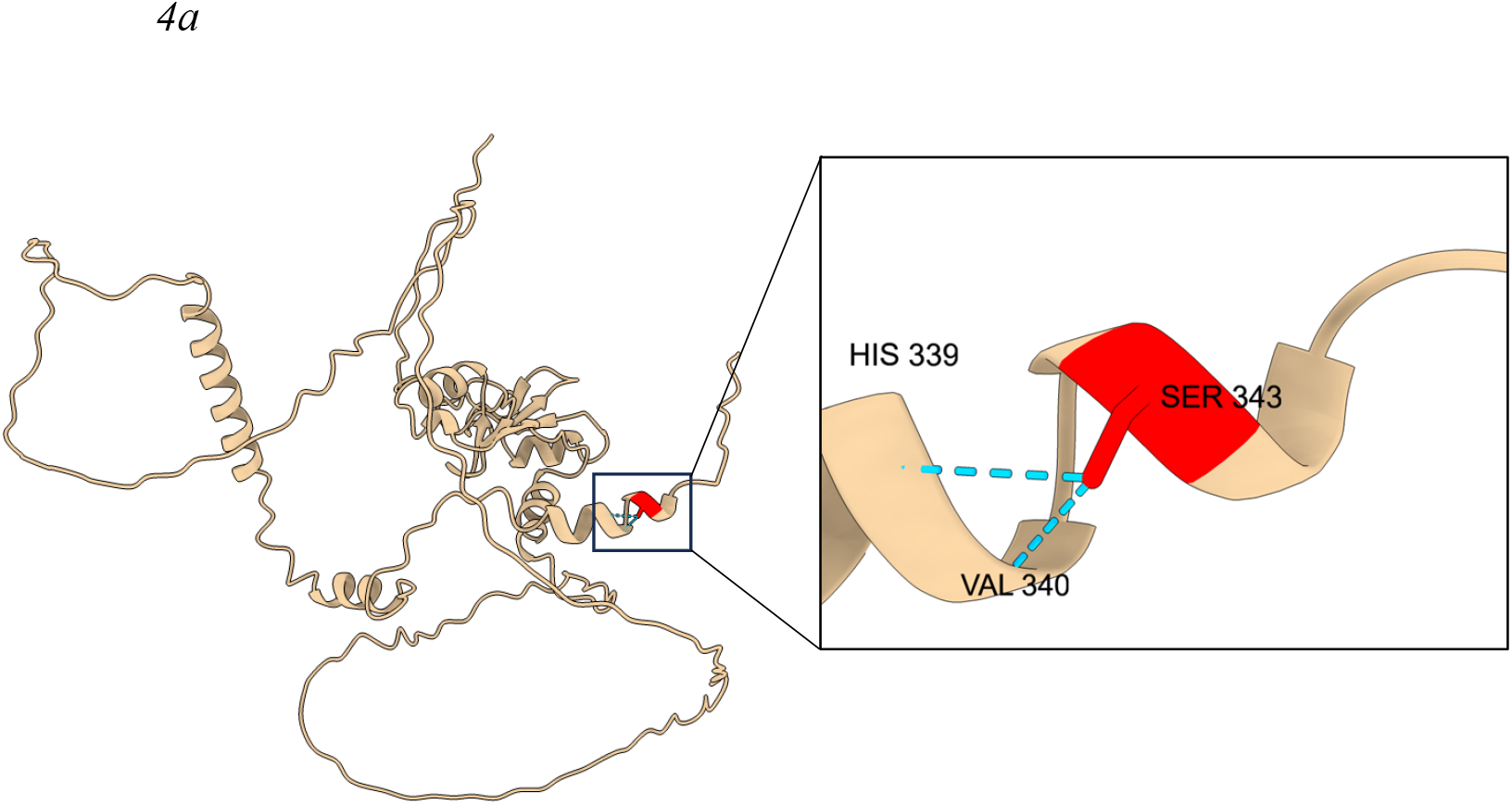
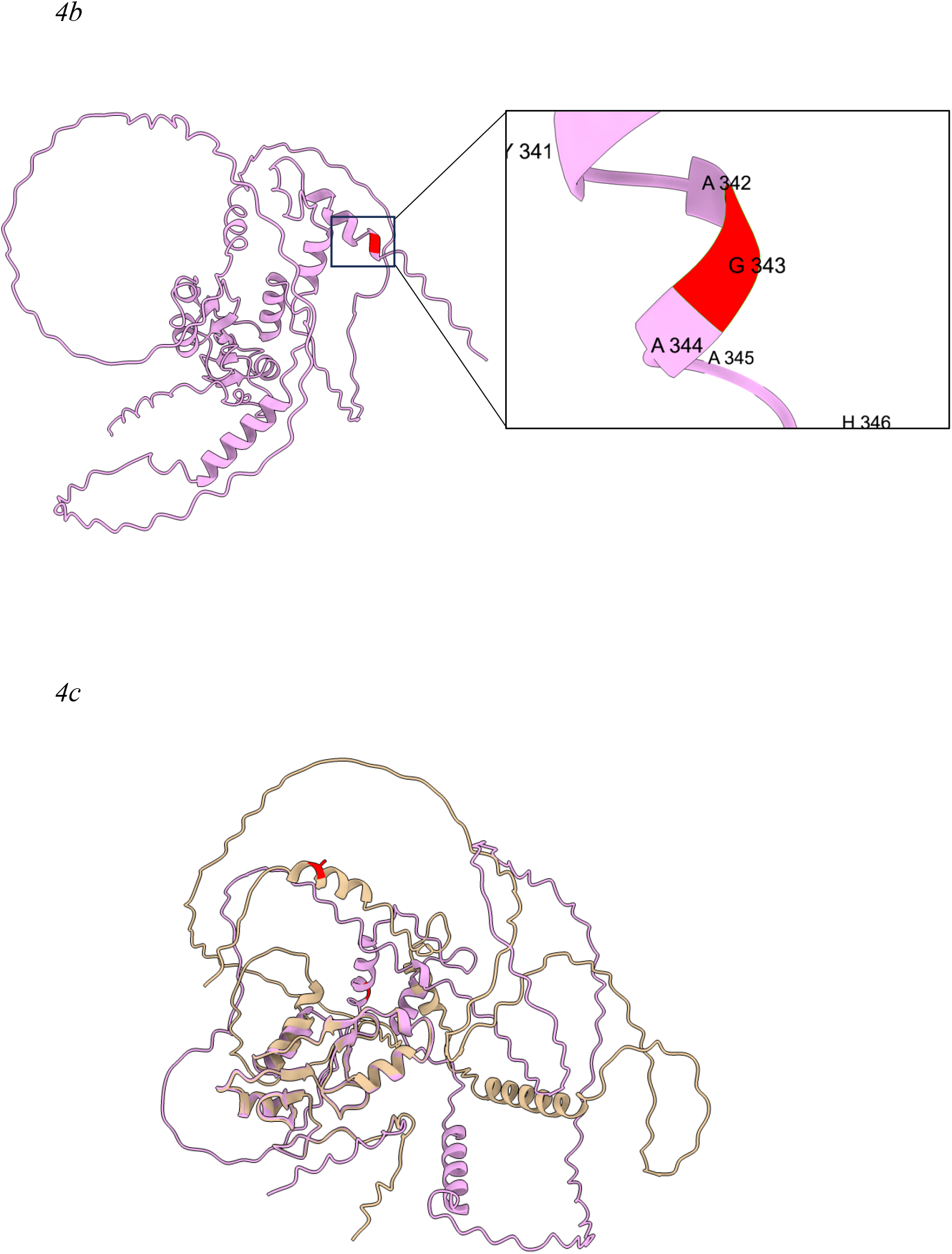
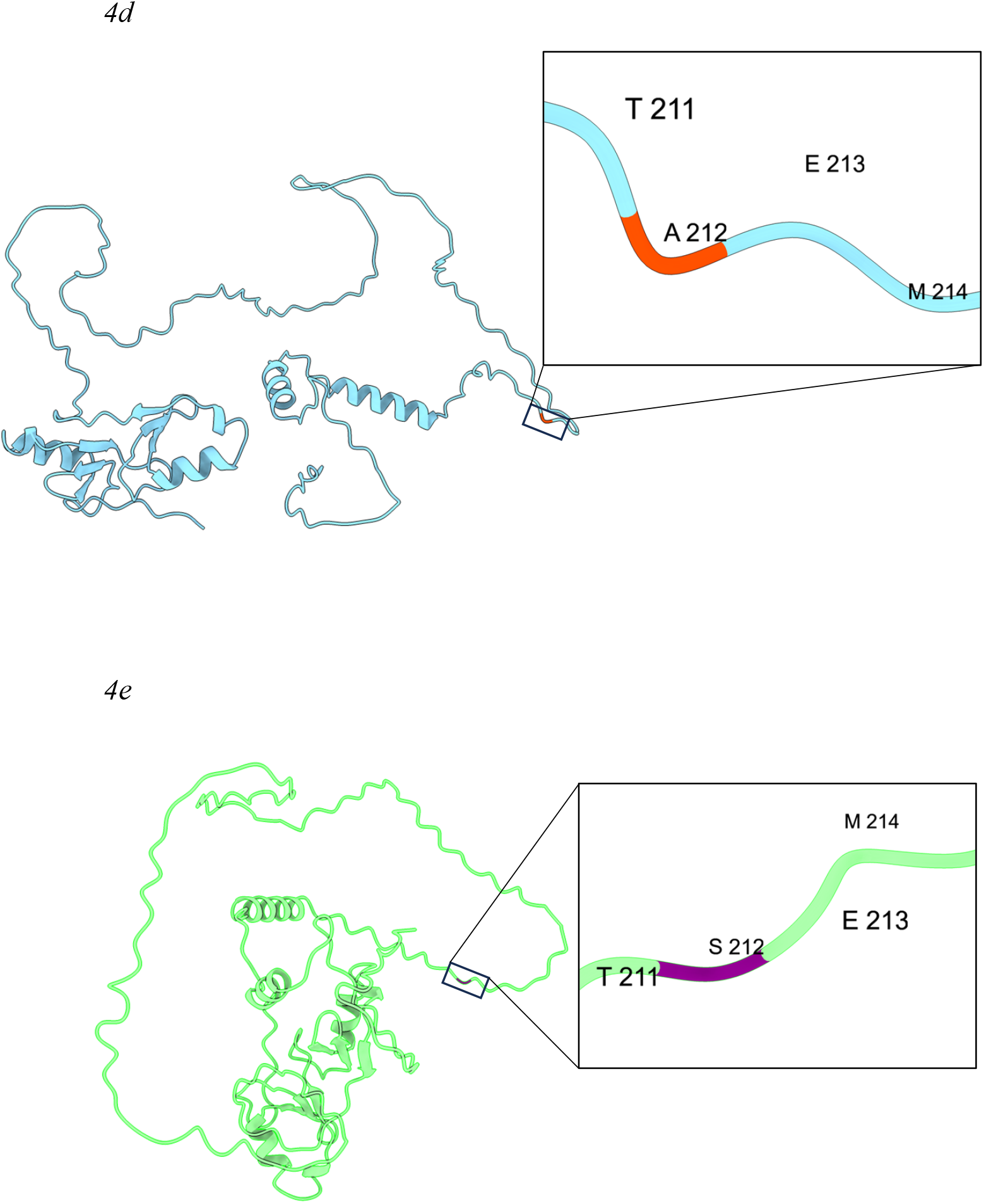
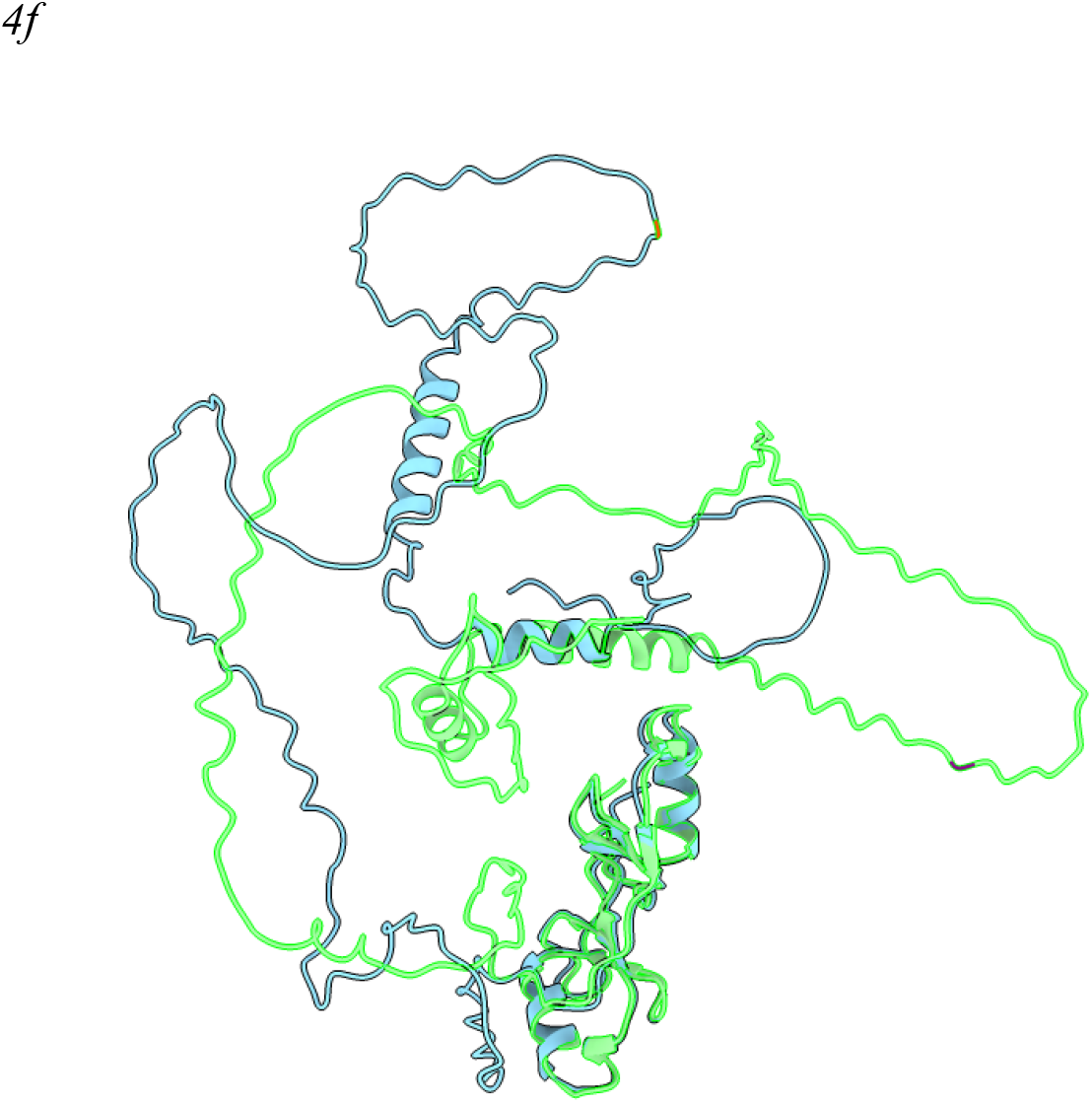
Structural impact of adaptive mutations in COL5 and COL3 proteins. (a) Ribbon representation of the wild-type COL5 protein with a close-up view of the S343 residue (red), highlighting its position within the structure. Hydrogen bonds involving S343 are indicated with sky-blue dashed lines. (b) Ribbon representation of the mutant COL5 (S343G) protein (red), arranged similarly to (a), showing the loss of hydrogen bonding at the mutation site. (c) Superimposition of wild-type COL5 (peach) and mutant G343S (light magenta) structures, revealing conformational changes. (d) Ribbon representation of the wild-type COL3 protein with a close-up view of the A212 (orange) residue, showing its position relative to the protein’s structural framework. (e) Ribbon representation of the mutant COL3 (A212A) protein (magenta), arranged similarly to (d), indicating no significant changes in hydrogen bonding. (f) Superimposition of wild-type COL3 (sky-blue) and mutant S212A (electric-green) structures, showing minimal overall conformational differences. Note structural models were generated using AlphaFold3 and visualized in ChimeraX. These representations highlight the distinct structural consequences of the adaptive substitutions, particularly the disruption of hydrogen bonding in COL5, which may affect protein stability and regulatory interactions. Also conformational changes for both COL3 and COL5 are visible in the superimposition images.

## 4. Discussion

This comprehensive analysis reveals that episodic positive selection has shaped key residues within the *Arabidopsis* COL gene family, particularly in COL3 and COL5, affecting primarily red and far-red light signaling as well as shoot and flower development, with profound implications for photoperiodic regulation.

Using CODEML-based site and branch-site models, statistically significant signals of adaptive evolution at specific codon sites were detected. The observation that some branches exhibit ω values significantly higher than the background (0.22) yet remain below 1 can be explained by a relative relaxation of purifying selection rather than strong positive selection^7^. In many genes, the majority of codon sites are under strong purifying selection, so even if a few sites experience episodic positive selection, the overall ω remains below 1^7^. This diluted signal may also reflect a transient or weak phase of adaptive evolution that does not overcome the dominant purifying forces acting on most sites.

Ancestral sequence reconstruction demonstrated that, for COL3, the ancestral state at residue 212 was serine, whereas the modern protein exhibits alanine. In COL5, the ancestral state at residue 343 was glycine, contrasting with the serine present in the modern protein. These shifts, though localized, suggest that episodic positive selection has driven functionally relevant substitutions that may modulate protein interactions and regulatory capacity.

GO enrichment analysis using ShinyGO revealed that the subset of COL genes exhibiting episodic positive selection is significantly enriched (BH-adjusted p < 0.05) for biological processes involved in red light signaling, regulation of flower development, regulation of shoot system development, and response to red or far-red light—with the red light signaling pathway showing the highest fold enrichment. In contrast, the remaining COL genes did not display significant enrichment for these pathways. This functional categorization supports the hypothesis that adaptive substitutions in COL genes are primarily associated with pathways critical for light perception and preferentially associated with pathways critical for developmental regulation, thereby enhancing plant responsiveness to environmental cues.

Structural modeling and stability analyses further support these findings. In COL5, the G343S adaptive mutation, which occurs outside the canonical CCT domain (285-327) within a unique proteomic region, was associated with modest stabilization and reduced flexibility, as indicated by DynaMut (ΔΔG = 0.195 kcal/mol) predictions, as well as a decrease in vibrational entropy (ΔΔSVib = -0.047 kcal·mol^−1^·K^−1^). Moreover, ChimeraX visualization revealed that the modern COL5 structure exhibits hydrogen bonding involving S343, which is lost when this residue is reverted to glycine, potentially altering regulatory interactions. In contrast, the COL3 S212A adaptive mutation, located immediately upstream of its CCT domain (229-271), exhibited a stabilizing effect based on DynaMut (ΔΔG = 0.735 kcal/mol) and resulted in a more pronounced reduction in flexibility (ΔΔSVib = –0.162 kcal·mol^−1^·K^−1^). Notably, no significant hydrogen bonding changes were observed in COL3 between wild-type and mutant forms, suggesting that the adaptive impact may be subtler, possibly affecting local conformational dynamics or protein-protein interaction surfaces which were evident from conformational analysis in ChimeraX.

Collectively, these results underscore that adaptive substitutions at specific sites can finely modulate the structural properties of COL proteins related to red and far-red light signaling pathways without overriding the strong purifying selection that governs the remainder of the gene. The alterations in protein stability and flexibility likely influence the functional performance of these COL proteins (COL3 and COL5) in mediating light signal transduction and photoperiodic flowering responses. By linking episodic positive selection to measurable changes in protein architecture, my findings provide a robust mechanistic framework for understanding how adaptive evolution contributes to the diversification of regulatory networks in plants. Future more in silico functional analysis will be essential to validate these structural predictions and to further elucidate the biological consequences of these adaptive modifications.

## 5. Conclusion

In conclusion, this study demonstrates that episodic positive selection has driven key adaptive substitutions in the COL gene family of Arabidopsis, notably affecting residues in COL3 and COL5 that are critical for light-responsive regulatory functions. Structural and stability analyses indicate that these mutations—reflected by measurable changes in free energy and protein flexibility—may fine-tune the interactions and conformational dynamics of COL proteins, thereby influencing photoperiodic regulation of flowering and plant development. Additionally, GO enrichment analysis revealed significant overrepresentation of red and farred light signaling pathways and developmental processes among the positively selected COL genes, reinforcing their adaptive significance. Future work should include molecular dynamics simulations to provide a more detailed understanding of the dynamic behavior of these proteins and experimental validations, such as site-directed mutagenesis and in vivo functional assays, to further delineate the biological consequences of these adaptive changes.

## 8. Data availability statement

The data are provided within the manuscript or supplementary information files. Also, the GitHub repository at https://github.com/sinhakrishnendu/AT_COL.git contains all the supplementary materials in the form of raw and processed data, as well as all necessary scripts to replicate the analyses presented in this manuscript.

## 9. Acknowledgments

I thank the global open-access community for their dedication to free science and for breaking down barriers to knowledge without which this study was not possible. I appreciate those who work to democratize information, driving progress and empowerment worldwide. I also acknowledge the freely available language models that helped refine this manuscript. Finally, I am deeply grateful to my wife, Dr. Nabanita Ghosh, Assistant Professor of Zoology at Maulana Azad College, Kolkata, for her insightful suggestions.

## 10. Declaration of generative AI and AI-assisted technologies in the writing process

During the preparation of this work the author(s) used ChatGPT in order to refine this manuscript’s readability. After using this tool/service, the author(s) reviewed and edited the content as needed and take(s) full responsibility for the content of the publication.

